# Comparative Neuroprotective Effects of Minocycline and Bone Marrow Mononuclear Cells After Complete Spinal Cord Transection in Adult Rats

**DOI:** 10.64898/2026.02.14.705644

**Authors:** Michelle Castro da Silva Holanda, Celice Cordeiro de Souza Bergh Pereira, Mário Santos Barbosa Junior, Jonabeto Vasconcelos Costa, Rosana Telma Lopes Afonso, Marcelo Marques Cardoso, Cláudio Eduardo Corrêa Teixeira, Edna Cristina Santos Franco, Walace Gomes Leal

## Abstract

Acute spinal cord injury triggers a complex secondary injury cascade characterized by lesion expansion, neuroinflammation, glial reactivity, and oligodendrocyte degeneration, which together limit endogenous repair. Identifying neuroprotective interventions capable of targeting distinct components of this cascade remains a major challenge. In this study, we compared the neuroprotective profiles of minocycline, a tetracycline derivative with anti-inflammatory and antioxidant properties, and bone marrow mononuclear cells (BMMCs), which exert paracrine immunomodulatory and trophic effects, using a model of complete thoracic spinal cord transection in adult rats. Animals received either BMMCs (5 × 10^6^ cells, intravenously, 24 h post-injury) or minocycline (50 mg/kg twice daily for 48 h, followed by 25 mg/kg for five days). Histological and immunohistochemical analyses revealed that both treatments attenuated secondary damage, reducing lesion area, microglial/macrophage activation (ED1^+^ cells), and oligodendrocyte pathology (Tau-1^+^ cells). However, the magnitude and pattern of protection differed between interventions: minocycline produced a stronger reduction in lesion area, whereas BMMCs exerted greater suppression of microglial/macrophage activation and superior preservation of oligodendrocytes. Astrocyte counts (GFAP^+^ cells) did not differ quantitatively among groups, despite qualitative differences in astrocytic morphology. Integrated effect size analysis further highlighted these complementary neuroprotective profiles across outcomes. Collectively, these findings indicate that minocycline and BMMCs target distinct components of secondary injury after severe spinal cord injury, providing a mechanistic rationale for future studies exploring multi-targeted or combinatorial therapeutic strategies.

## Introduction

Acute spinal cord injury triggers a secondary injury cascade that amplifies the initial trauma and remains a barrier to functional recovery [1, 2]. For decades, research has focused on developing neuroprotective strategies to mitigate this damage, with two major avenues being pharmacological agents and cell-based therapies. Among these, the antibiotic minocycline and bone marrow-derived mononuclear cells (BMMCs) have emerged as leading candidates, backed by extensive preclinical evidence demonstrating their ability to modulate inflammation and promote tissue preservation [3, 4].

However, the translation of these promising preclinical findings into effective clinical treatments has been challenging. Despite their individual strengths, both minocycline and BMMC-based therapies have yielded inconclusive results in human clinical trials when used as monotherapies [5]. This translational gap has prompted a critical paradigm shift within the spinal cord injury field, away from the search for a single therapy approach and toward the development of multi-targeted combination therapies designed to simultaneously address multiple facets of the complex secondary injury cascade [6].

It is in this context that a comparison of minocycline and BMMCs becomes relevant. While both are broadly termed neuroprotective, their underlying mechanisms are fundamentally different. Minocycline acts as a broad-spectrum inhibitor of microglial activation and apoptosis [7], while BMMCs are thought to function primarily through sophisticated paracrine signaling, orchestrating a complex immunomodulatory and trophic response [8]. To date, however, these two distinct therapeutic modalities have not been comparatively evaluated within the same severe spinal cord injury model to elucidate their unique and potentially complementary contributions to neuroprotection.

In a previous investigation, we have compared the therapeutic effects of both minocycline and BMMCs following striatal stroke, rendering important peculiarities for both therapeutic approaches [9, 10]. So far, such a comparative investigation had not been performed. Therefore, the present study was designed to compare the distinct neuroprotective profiles of systemic minocycline administration and intravenous BMMC transplantation. By quantifying their differential impacts on lesion size, microglial/macrophage activation, astrogliosis, and oligodendrocyte preservation, we aim to provide a clear, mechanistic rationale for how these two therapies could be combined in a future, more effective, multi-targeted strategy for treating acute spinal cord injury.

## Methods

### Experimental Animals

Sixteen adults male Wistar rats (n = 16), weighing between 250 and 300 g, were used and maintained under controlled laboratory conditions. All procedures followed national guidelines for animal experimentation and were approved by the Animal Ethics Committee of the Federal University of Pará (protocol no. BIO001-2011). One animal was the donor of BMMCs cells. All other animals were allocated into four experimental groups: sham operated group (n = 3), which underwent laminectomy without spinal cord injury; control group (n = 4), submitted to complete transection of the spinal cord and treated only with sterile saline; BMMC group (n = 4), submitted to the same lesion and treated 24 hours after trauma with 5 × 10^6^ BMMCs administered intravenously; and the minocycline (MIN) group (n = 4) treated with minocycline administered intraperitoneally for seven consecutive days. One additional animal from the same litter was used exclusively as a donor for the isolation of BMMCs cells.

### Surgical Procedure

To induce spinal cord lesion, animals were anesthetized with ketamine hydrochloride (72 mg/kg) and xylazine (9 mg/kg), administered intraperitoneally. After abolition of corneal and hindlimb withdrawal reflexes, dorsal trichotomy was performed, followed by placement in a stereotaxic apparatus and asepsis of the surgical area. A 4-cm incision allowed exposure of the paravertebral musculature, which was retracted to visualize the vertebral lamina at T8.

A total laminectomy was performed using a micro-rongeur, exposing the spinal cord, and complete transection was carried out with a scalpel and confirmed by surgical microscopy. A complete transection is not very common in the clinics, but such model was chosen in order to investigate the treatment efficacy in an extreme experimental condition. After this procedure, skin was sutured, and animals were housed individually, monitored for hydration, food intake, and excretory function, and received bladder expression twice daily. During the first five postoperative days, all animals received antibiotic treatment with 10% enrofloxacin (2.5 mg/kg, intramuscular).

### Treatment with Bone Marrow Mononuclear Cells

Mononuclear cells were obtained from the donor animal, anesthetized with the same protocol and euthanized by cervical dislocation. The tibias and femurs were dissected and kept in saline until processing in a laminar flow hood. The bone epiphyses were removed, and the bone marrow was flushed with DMEM-F12, dissociated, and centrifuged for 5 minutes. After an additional dissociation, the material was subjected to a Histopaque® gradient and centrifuged for 30 minutes. The mononuclear fraction was collected, washed three times in sterile saline, and resuspended in DMEM-F12 supplemented with fetal bovine serum. Viable cells were counted in a Neubauer chamber after mixing with Trypan Blue, and the final suspension volume was adjusted to obtain 5 × 10^6^ cells per animal in the BMMC group.

### Minocycline Treatment

Minocycline administration followed the protocol published in [9]. In brief, animals received two daily intraperitoneal doses of 50 mg/kg during the first 48 hours post-injury, with a 12-hour interval between doses, followed by a single daily dose of 25 mg/kg for the subsequent five days.

### Histopathological Analysis

Seven days after lesion induction, all animals were re-anesthetized with ketamine and xylazine and perfused via the left cardiac ventricle with 500 mL of heparinized saline, followed by 500 mL of 4% paraformaldehyde. Spinal cords were removed, post-fixed for 24 hours, and cryoprotected. The lesion region was embedded in Tissue-Tek® OCT and longitudinal sections of 20 µm and 50 µm were obtained in a cryostat. Slides were stored at –20 °C until processing. The 50-µm sections were stained with cresyl violet (Nissl) for qualitative assessment and estimation of lesion area, which was later analyzed using ImageJ software. The 20-µm sections were used for immunohistochemistry.

### Immunohistochemistry

Immunohistochemical analyses used the antibodies anti-Tau-1 (1:500, a marker of pathological oligodendrocytes), anti-GFAP (1:1000, a marker of astrocytes), and anti-ED-1 (1:200, a marker of activated microglia/macrophages). Slides were thawed, washed in PBS, and subjected to heat treatment in borate buffer at 65 °C, followed by peroxidase blocking with a methanol–hydrogen peroxide mixture. Sections were then incubated with normal horse serum (for ED-1 and Tau-1) or goat serum (for GFAP), followed by overnight incubation with primary antibodies. The following day, slides were incubated with appropriate biotinylated secondary antibodies, treated with avidin–biotin complex, and developed with DAB. After dehydration and clearing, sections were mounted on permanent slides.

### Data Analysis

Qualitative descriptive analysis was performed under a light microscope with digital image capture. Quantitative analyses were performed using summarized data reported as mean ± standard deviation (s.d.), with the number of animals (*n*) per group. Individual-level data were not available; therefore, analyses were restricted to methods appropriate for aggregated datasets and small sample sizes. Lesion area and ED1+, GFAP+, and Tau-1+ cell counts were evaluated in the sham, control, BMMC and MIN groups.

Pairwise comparisons were defined *a priori*: control vs sham (model validation), BMMC vs control and MIN vs control (treatment effects), and BMMC vs MIN (secondary comparison). Effect sizes were estimated using Hedges’ *g*, which corrects for small-sample bias, with corresponding 95% confidence intervals (95% CI) used to express uncertainty and enable comparison across outcomes. Group differences were characterized using effect size estimation (Hedges’ *g*) with 95% confidence intervals. Emphasis was placed on estimation rather than null-hypothesis significance testing due to the small sample size.

Results are presented using representative histopathological images, summary tables, and a forest plot displaying effect sizes and 95% CI. All analyses were performed in Python. Given the absence of individual-level data, results are interpreted as estimation-focused and exploratory.

## Results

### Experimental model and general observations

All animals submitted to complete spinal cord transection developed immediate flaccid paralysis of the hind limbs, absence of reflex responses, and neurogenic bladder, confirming the effectiveness and reproducibility of the injury model. Sham-operated animals did not exhibit neurological deficits.

### Lesion area

Cresyl violet staining revealed preserved cytoarchitecture in sham-operated animals, with clear distinction between white and gray matter (Figure 1A). In contrast, untreated control animals exhibited extensive necrotic areas at the lesion site, characterized by tissue disorganization and cavitation (Figure 1B). Animals treated with MIN (Figure 1C) or BMMCs (Figure 1D) showed visibly reduced lesion areas compared with untreated control animals.

**Figure 1.**
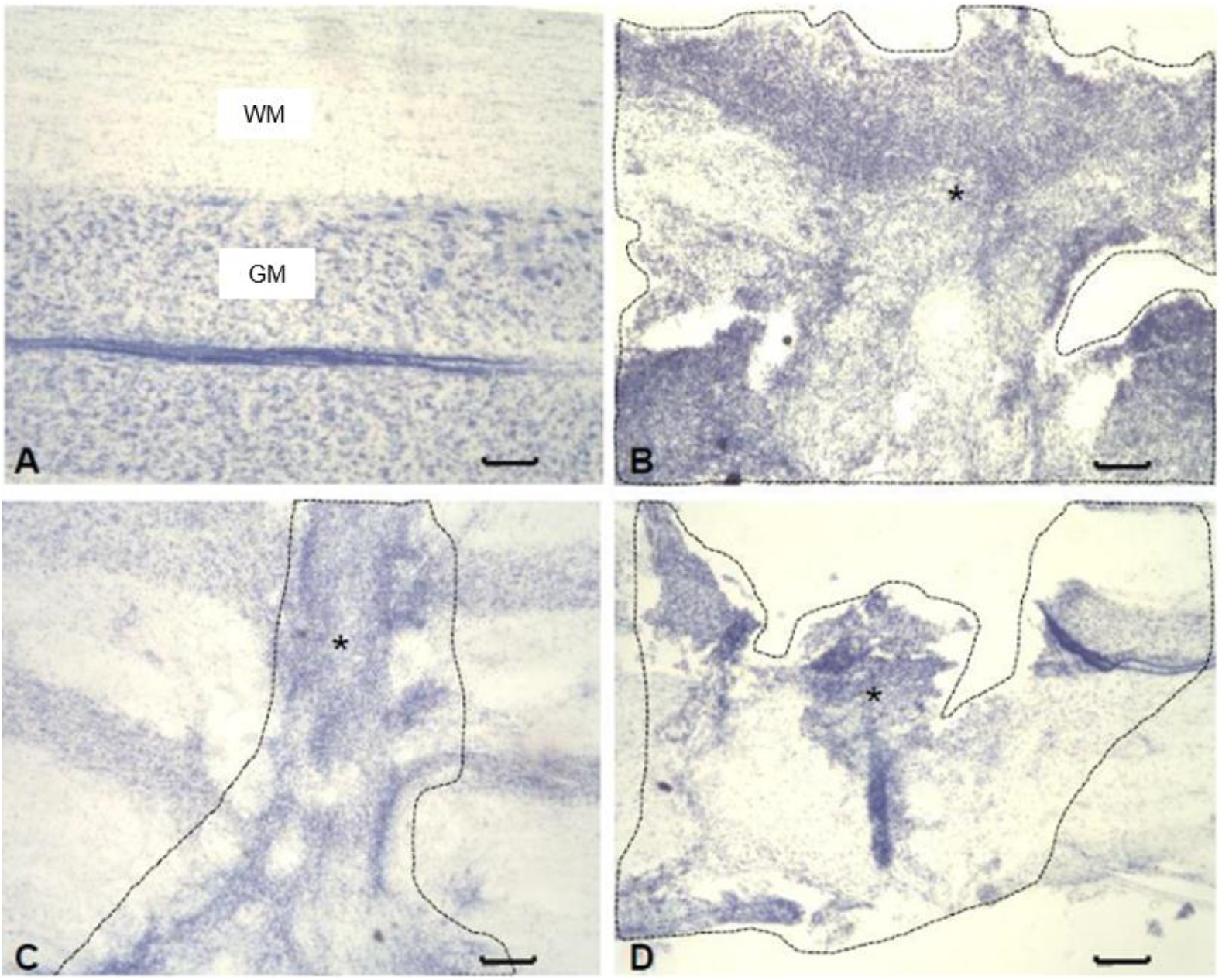
Necrotic lesion area after spinal cord transection visualized by cresyl violet staining. (A) Sham; (B) Control; (C) MIN-treated; (D) BMMCs–treated. Dashed lines indicate the necrotic lesion area used for quantification. WM, white matter; GM, gray matter. Asterisks (*) mark the lesion center. Scale bar: 200 μm.

Quantitative analysis revealed a marked reduction in lesion area in treated animals compared to control ones (Figure 2). Effect size estimation confirmed large negative effects for both BMMCs vs control (Hedges’ *g* = −8.52; 95% CI: −14.01 to −3.03) and MIN vs control (Hedges’ *g* = −12.72; 95% CI: −21.68 to −3.76), indicating substantial attenuation of tissue damage. Direct comparison between treatments revealed a positive effect size favoring MIN (BMMCs vs MIN Hedges’ *g* = 6.32; 95% CI: 2.13 to 10.52).

**Figure 2.**
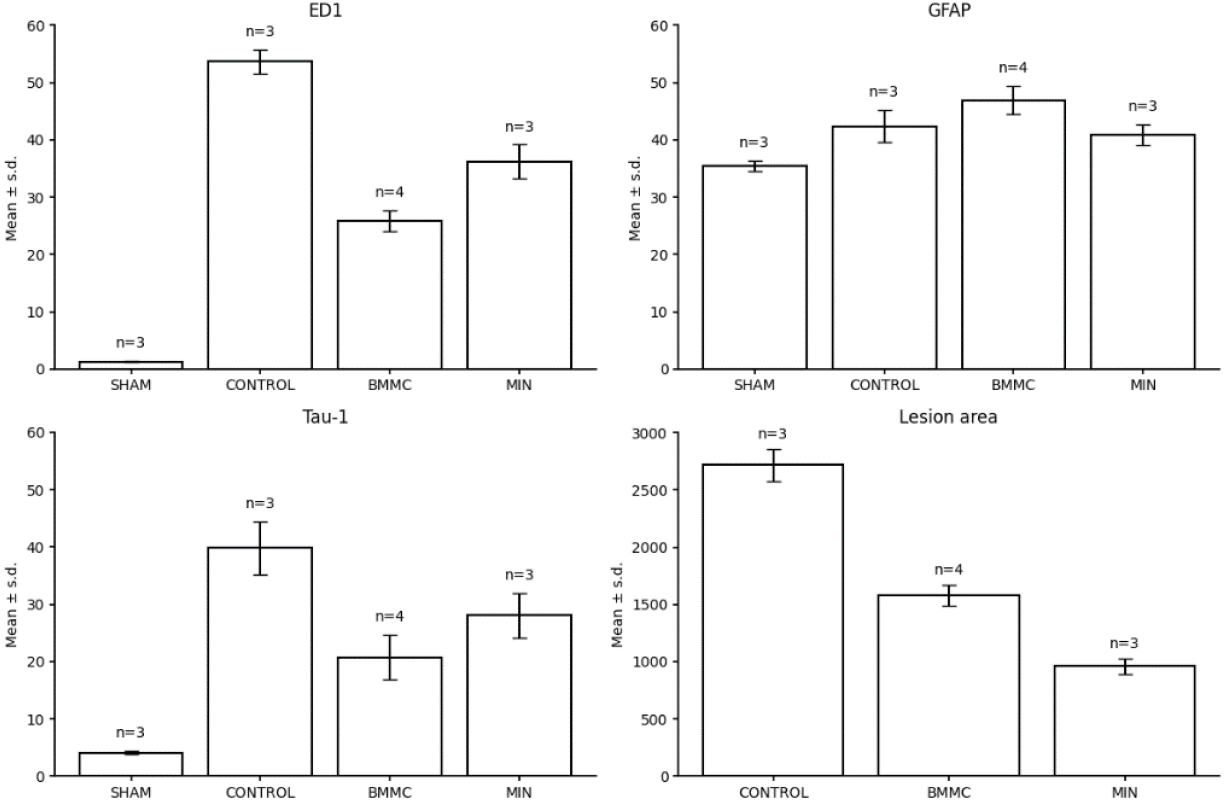
Values are expressed as Mean ± sd for lesion area and ED1+, GFAP+, and Tau-1+ cell counts per field in the sham, control, BMMC and MIN groups. *n* denotes the number of animals per group.

### Macrophage/microglial activation (ED1+ cells)

Immunohistochemical analysis using the anti-ED1 antibody has shown minimal macrophage/microglial activation in sham-operated animals (not illustrated). In untreated transected animals, a marked accumulation of ED1^+^ cells were observed at the lesion site (Figure 3A-B). Treatment with BMMCs (Figure 3C-D) or MIN (Figure 3E-F) visibly reduced ED1^+^ cell density.

**Figure 3.**
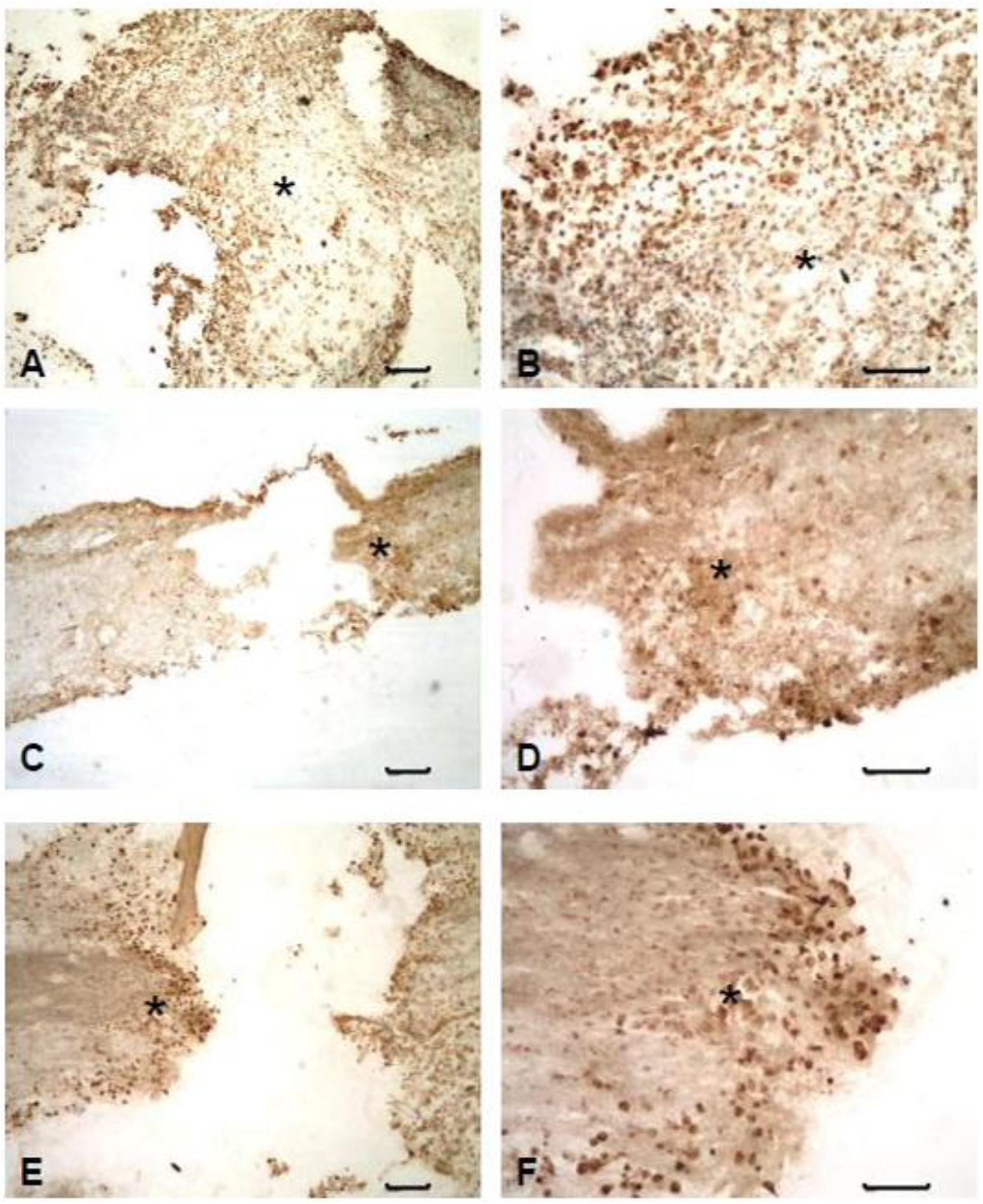
Macrophage/microglial activation detected by anti-ED1 immunohistochemistry. (A, B) Control; (C, D) BMMCs–treated; (E, F) MIN-treated. Panels A, C, and E show low magnification (200 μm), and B, D, and F high magnification (50 μm). Asterisks (*) indicate the lesion center.

Quantification confirmed a significant increase in ED1+ cells in the control group compared with sham (Figure 2). Effect size analysis showed an extremely large positive effect for control vs sham (Hedges’ *g* = 29.05; 95% CI: 8.86 to 49.24), validating the inflammatory response induced by injury. Both treatments reduced ED1+ cell counts relative to control, with a larger magnitude for BMMCs vs control (Hedges’ *g* = −12.40; 95% CI: −20.23 to −4.57) compared with MIN vs control (Hedges’ *g* = −5.50; 95% CI: −9.64 to −1.37). The direct comparison (BMMCs vs MIN) further favored BMMCs (Hedges’ *g* = −3.74; 95% CI: −6.50 to −0.98).

### Astrocytic response (GFAP+ cells)

GFAP immunohistochemistry revealed astrocytic labeling in all experimental groups (Figures 4-5). Sham-operated animals exhibited a typical astrocytic network with preserved morphology. In injured animals, astrocytes displayed hypertrophy and process thickening, particularly in the perilesional region, regardless of treatment.

**Figure 4.**
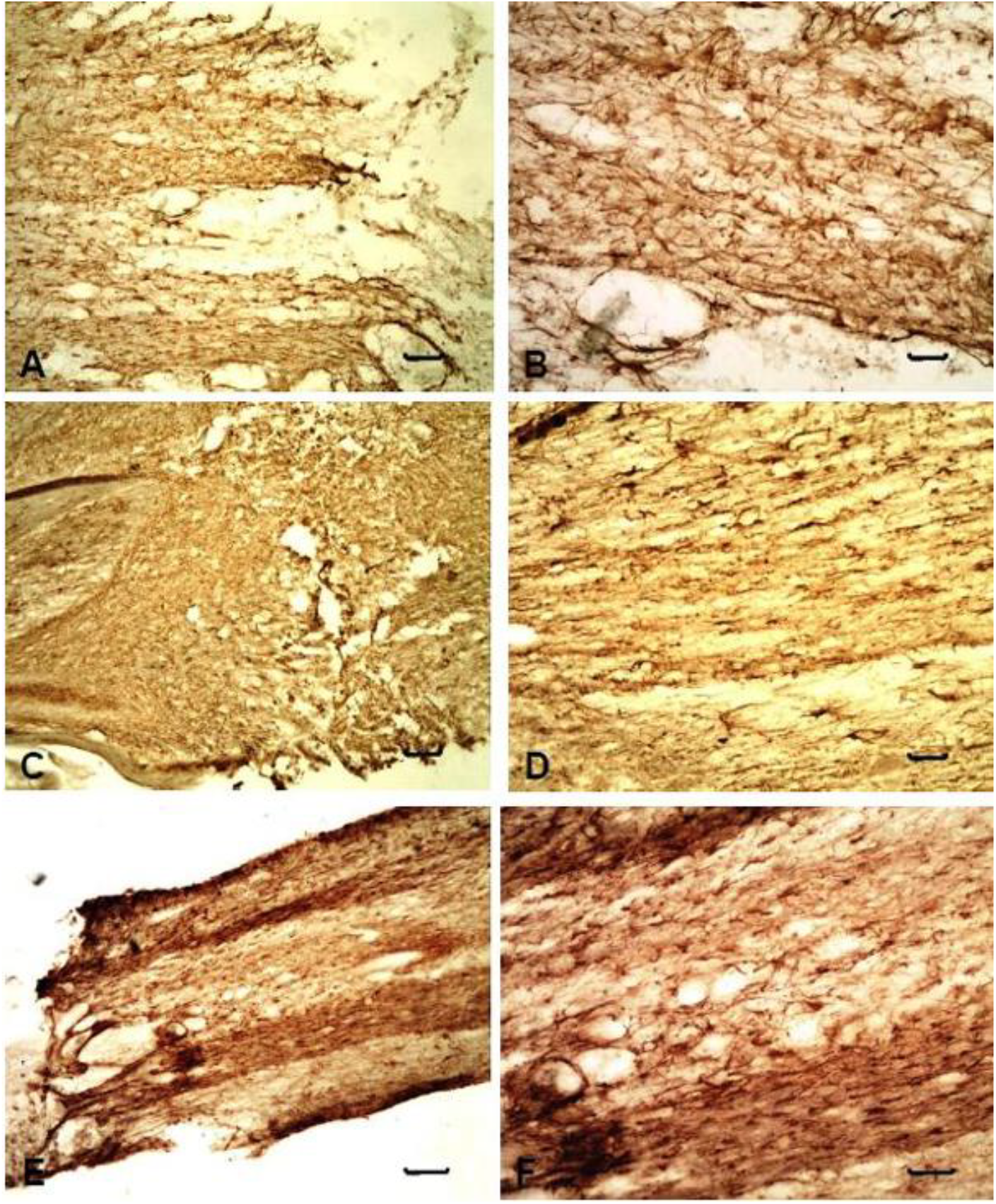
Astrocyte activation assessed by immunohistochemistry using an anti-GFAP antibody. (A, B) Control; (C, D) BMMCs-treated; (E, F) MIN-treated. Panels A, C, and E show low-magnification views (scale bar: 200 μm), whereas panels B, D, and F show higher-magnification views (scale bar: 50 μm).

**Figure 5.**
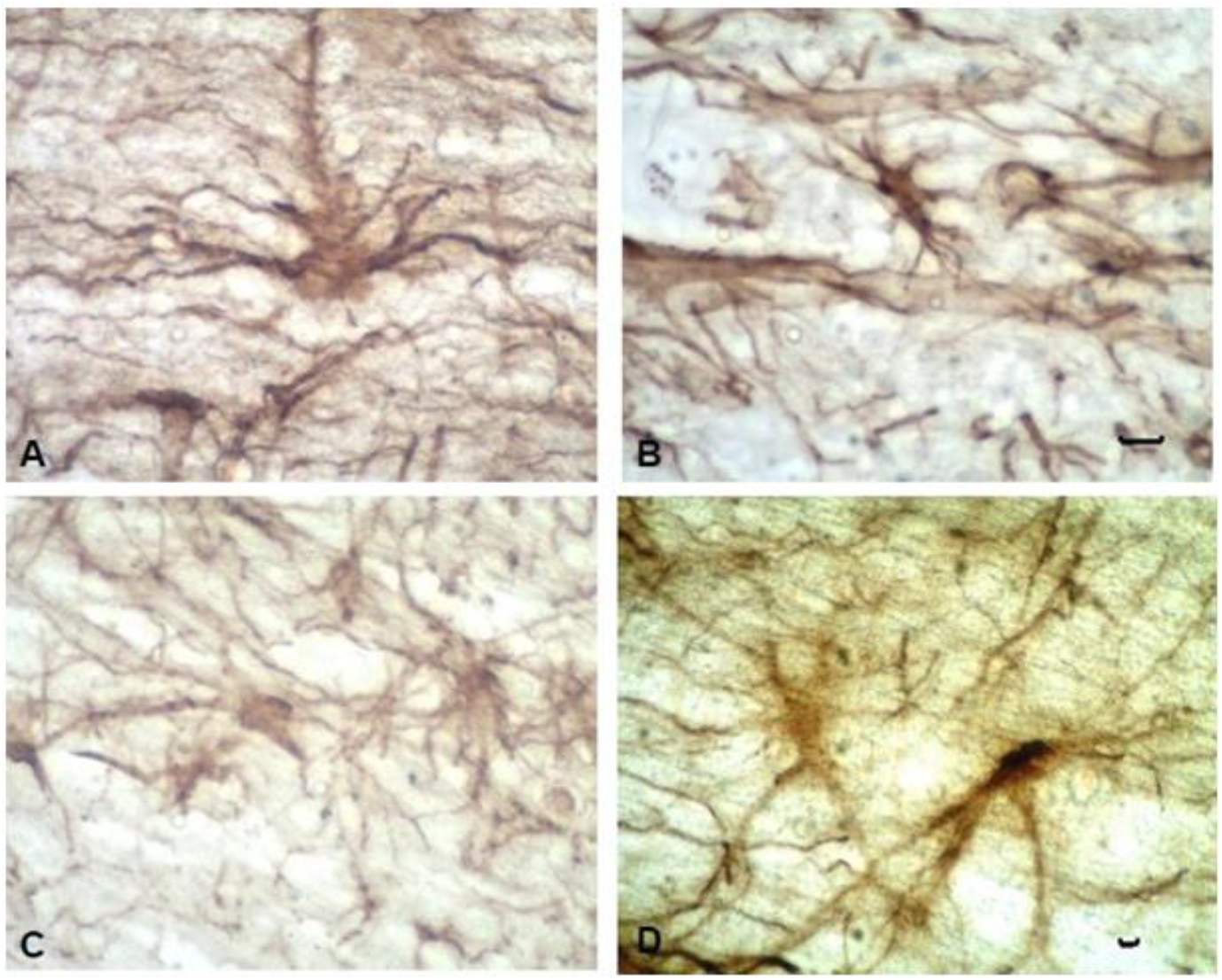
Astrocytic reactivity in the different experimental groups. (A) Sham; (B) control; (C) BMMCs– treated; (D) MIN-treated animal. Images were acquired using a 40× objective (scale bar: 50 μm) and illustrate white matter of the spinal cord.

Quantitative analysis showed modest differences in GFAP+ cell counts across groups (Figure 2). Effect size estimates indicated a small-to-moderate increase for control vs sham animals (Hedges’ *g* = 2.69; 95% CI: 0.24 to 5.15). Comparisons involving treatments yielded small or overlapping confidence intervals (BMMCs vs control, MIN vs control), whereas the direct comparison between BMMCs vs MIN suggested a moderate effect favoring BMMCs (Hedges’ *g* = 2.28; 95% CI: 0.22 to 4.34). Overall, these findings indicate that astrocytic changes were predominantly morphological rather than numerical.

### Oligodendrocyte pathology (Tau-1+ cells)

Tau-1 immunoreactivity was minimal in sham-operated animals (not illustrated). Untreated control animals exhibited a marked increase in Tau-1+ cells at the lesion site, consistent with oligodendrocyte pathology (Figure 6A). Both BMMCs (Figure 6B) and MIN (Figure 6C) treatments reduced Tau-1 immunoreactivity.

**Figure 6.**
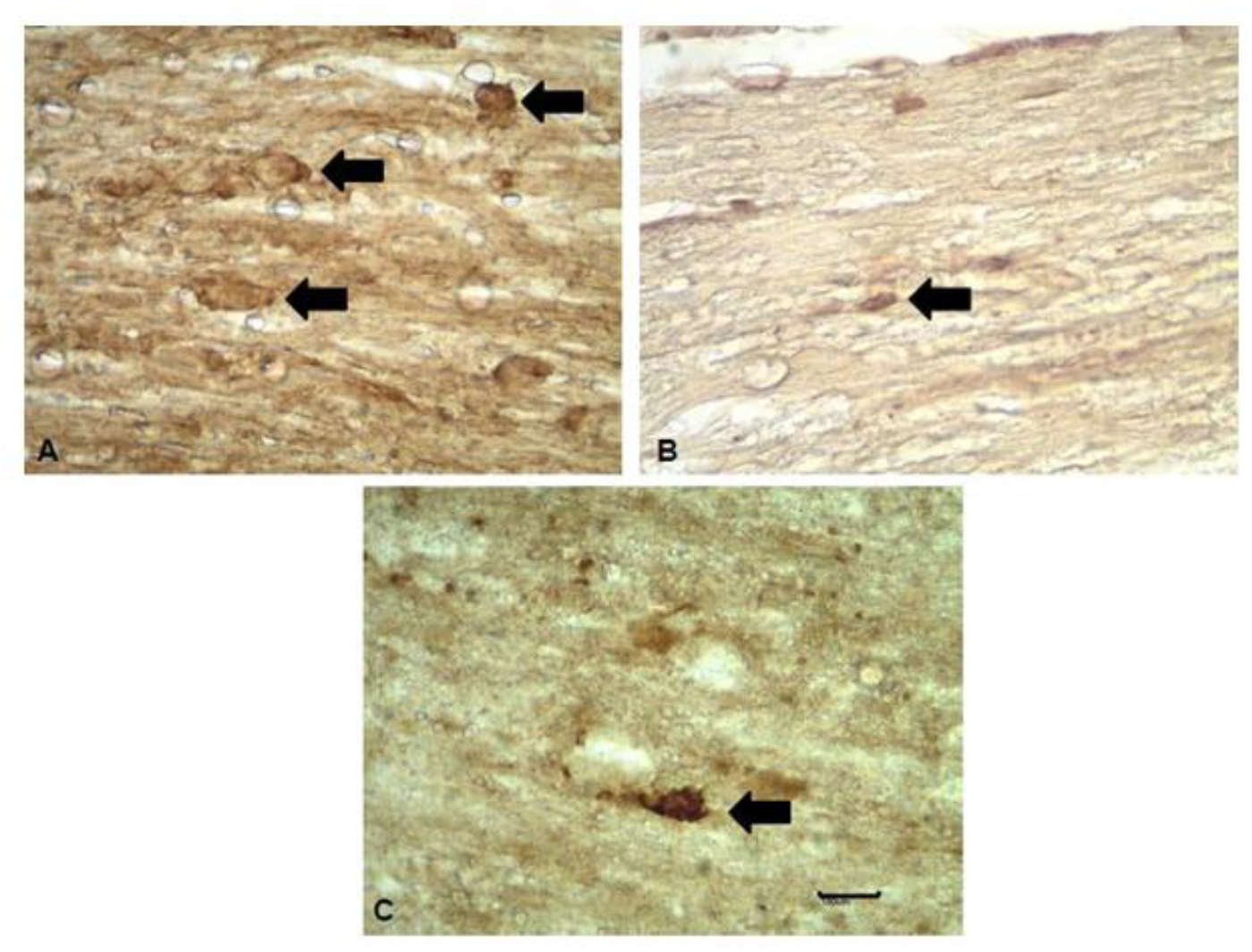
Tau-1 expression following complete spinal cord transection and treatment with MIN or BMMCs. Control; (B) BMMC-treated; (C) MIN-treated. Arrows indicate Tau-1–positive cells. Scale bar: 100 μm.

Quantitative analysis confirmed a substantial increase in Tau-1+ cells in the control group compared with sham (Figure 2). Effect size estimates demonstrated a large positive effect for control vs sham (Hedges’ *g* = 8.83; 95% CI: 2.51 to 15.16). Treatment effects were more pronounced for BMMCs vs control (Hedges’ *g* = −3.85; 95% CI: −6.66 to −1.03) than for MIN vs control (Hedges’ *g* = −2.21; 95% CI: −4.43 to 0.00). Comparisons between treatments (BMMCs vs MIN) favored BMMCs (Hedges’ *g* = −1.57; 95% CI: −3.35 to 0.22). BMMCs exhibited greater effects on inflammatory modulation and oligodendrocyte pathology.

### Integrated effect size analysis

To integrate the magnitude and direction of treatment effects across outcomes, effect size estimates for all planned comparisons are summarized in Table 1 and illustrated in the forest plot (Figure 7). This analysis highlights complementary treatment profiles, with minocycline showing a stronger effect on lesion area reduction, whereas BMMC exhibited greater effects on inflammatory modulation and oligodendrocyte pathology.

**Table 1.** Effect size estimates for histopathological outcomes following spinal cord transection.

| **Outcome** | **Comparison** | **Hedges' g** | **CI 95% (low)** | **CI 95% (high)** |
| --- | --- | --- | --- | --- |
| **ED1** | **CONTROL vs SHAM** | 29.05 | 8.86 | 49.24 |
| **ED1** | **BMMC vs CONTROL** | -12.4 | -20.23 | -4.57 |
| **ED1** | **MIN vs CONTROL** | -5.5 | -9.64 | -1.37 |
| **ED1** | **BMMC vs MIN** | -3.74 | -6.5 | -0.98 |
| **GFAP** | **CONTROL vs SHAM** | 2.69 | 0.24 | 5.15 |
| **GFAP** | **BMMC vs CONTROL** | 1.47 | -0.28 | 3.22 |
| **GFAP** | **MIN vs CONTROL** | -0.52 | -2.16 | 1.12 |
| **GFAP** | **BMMC vs MIN** | 2.28 | 0.22 | 4.34 |
| **Tau-1** | **CONTROL vs SHAM** | 8.83 | 2.51 | 15.16 |
| **Tau-1** | **BMMC vs CONTROL** | -3.85 | -6.66 | -1.03 |
| **Tau-1** | **MIN vs CONTROL** | -2.21 | -4.43 | 0.0 |
| **Tau-1** | **BMMC vs MIN** | -1.57 | -3.35 | 0.22 |
| **Lesion area** | **BMMC vs CONTROL** | -8.52 | -14.01 | -3.03 |
| **Lesion area** | **MIN vs CONTROL** | -12.72 | -21.68 | -3.76 |
| **Lesion area** | **BMMC vs MIN** | 6.32 | 2.13 | 10.52 |

Effect sizes are reported as Hedges' $g$ with 95% confidence intervals for pre-specified comparisons among the SHAM, CONTROL, BMMC, and MIN groups. Positive $g$ values indicate higher outcome values in the first group of each comparison, and negative values indicate lower values. Estimates were corrected for small-sample bias, allowing comparison of effect magnitude and direction across histopathological outcomes.

**Figure 7.**
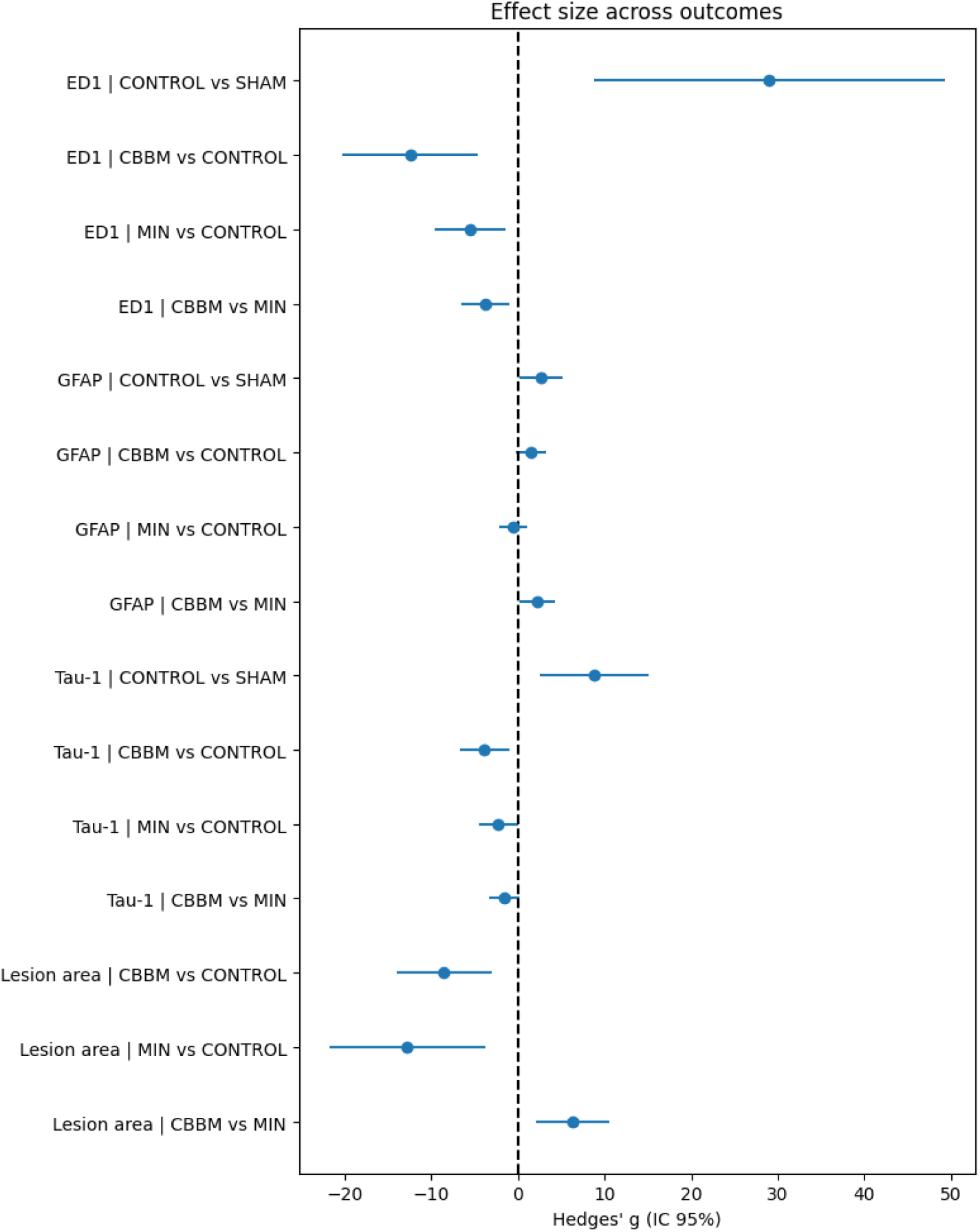
Forest plot of effect sizes (Hedges’ *g*, 95% CI) for histopathological outcomes after spinal cord transection, showing pre-specified comparisons among SHAM, CONTROL, CBBM, and MIN groups. Positive *g* values indicate higher outcome values in the first group, and negative values indicate lower values. The dashed line at zero denotes no effect.

## Discussion

The present study was designed to comparatively evaluate the neuroprotective profiles of minocycline and BMMCs in a severe model of complete spinal cord transection. By exa--mining lesion size, inflammatory activation, astrocytic response, and oligodendrocyte pathology within the same experimental framework, we aimed to clarify whether these two widely studied interventions exert overlapping or distinct biological effects. Our findings demonstrate that, although both treatments confer histopathological protection, they do so through distinct and complementary mechanisms, supporting the rationale for multi-targeted therapeutic strategies in acute spinal cord injury. One of the most prominent findings was the differential effect on lesion area. Minocycline produced a greater reduction in lesion size compared with BMMC treatment, consistent with its pleiotropic pharmacological actions, including inhibition of microglial activation, suppression of apoptotic pathways, and attenuation of oxidative stress [3,7]. These mechanisms are particularly relevant during the acute phase of SCI, when secondary injury processes rapidly expand the initial lesion [1,2]. Despite this robust tissue-sparing effect, clinical translation of minocycline has remained limited. Recent systematic reviews indicate that, although biologically active, minocycline has not consistently produced meaningful neurological improvement in human spinal cord injury when administered as a monotherapy [5]. Our results reinforce the notion that containment of lesion expansion alone may be insufficient to counteract the multifactorial nature of secondary injury.

In contrast, BMMC treatment exerted a more pronounced effect on the inflammatory response, as evidenced by a stronger reduction in ED1^+^ macrophage/microglial activation. This finding highlights the immunomodulatory potential of bone marrow–derived cell populations, which act predominantly through paracrine mechanisms rather than direct cell replacement [4,8]. The ability of BMMCs to reshape the inflammatory milieu toward a more reparative profile aligns with growing evidence that targeted modulation of neuroinflammation, rather than global suppression, is critical for promoting tissue preservation and repair after spinal cord injury [11, 12]. Importantly, excessive or prolonged microglial activation is a major contributor to secondary degeneration, yet microglia also play essential roles in debris clearance and coordination of repair processes. The more nuanced immunomodulation observed with BMMCs may therefore represent a therapeutic advantage over broadly acting anti-inflammatory agents.

Preservation of oligodendrocytes emerged as another key distinction between the two interventions. BMMC treatment was associated with a greater reduction in Tau-1^+^ pathological oligodendrocytes compared with minocycline. This observation is particularly relevant in light of the evolving understanding that long-term functional recovery depends not only on axonal survival but also on the maintenance of myelinating cells and white matter integrity [2]. Protection of oligodendrocytes is essential for preserving axonal conductivity in spared fibers and for supporting endogenous repair mechanisms. The superior oligodendrocyte-sparing effect of BMMCs suggests that cell-based therapies may address aspects of spinal cord injury pathology that are less effectively targeted by pharmacological agents alone.

Astrocytic responses, assessed by GFAP immunoreactivity, exhibited relatively modest quantitative differences among groups, despite clear qualitative changes in astrocyte morphology. Effect size estimates indicated that treatment-related effects on astrocytes were predominantly morphological rather than numerical. This finding is consistent with the dynamic role of astrocytes after spinal cord injury, where changes in cellular architecture, process hypertrophy, and spatial organization may be more informative than absolute cell counts.

Taken together, integration of effect size estimates across all histopathological outcomes reveals a coherent pattern: minocycline is more effective in limiting lesion expansion, whereas BMMCs exert stronger effects on inflammatory modulation and oligodendrocyte preservation. These complementary profiles provide mechanistic insight into why single-agent neuroprotective strategies have yielded limited success in clinical trials and support the growing consensus that effective spinal cord injury therapies must simultaneously target multiple components of the secondary injury cascade [6].

Several limitations of the present study should be acknowledged. The analyses were based on summarized data and small sample sizes, precluding individual-level inference and multivariate analyses. Accordingly, emphasis was placed on estimation of effect magnitude and confidence intervals rather than exclusive reliance on null-hypothesis testing. Despite these constraints, the consistency of treatment effects across independent histopathological measures strengthens the biological relevance of the findings.

## Conclusion

This study demonstrates that minocycline and bone marrow mononuclear cells confer neuroprotection after complete spinal cord transection through distinct and complementary mechanisms. Minocycline preferentially attenuates lesion expansion, whereas BMMCs exert stronger immunomodulatory and oligodendrocyte-preserving effects. These findings provide a mechanistic framework that helps explain the limited efficacy of monotherapies observed in clinical trials and support the development of multi-targeted or combinatorial therapeutic strategies for acute spinal cord injury.

